# Multi-Omics Insights into Grapevine Ecodormancy to Budburst Transition: Interplay of Gene Expression, miRNA Regulation, and DNA Methylation

**DOI:** 10.1101/2023.10.21.563414

**Authors:** Harshraj Shinde, Tajbir Raihan, Lakshay Anand, Sharyn E. Perry, Robert L. Geneve, Carlos M. Rodriguez Lopez

## Abstract

In grapevine, the transition from a dormant bud to budburst is a critical developmental process related to vegetative and reproductive growth. We generated a time series analysis (five sampling time points) and used transcriptome, small RNA, and whole-genome bisulfite sequencing to characterize this transition. Ecodormant buds took an average of 17 days to budburst. Transcriptome analysis identified a total of 7002 differentially expressed genes across all sampling times and revealed that the brassinosteroid metabolism and the linoleic acid metabolism pathways are upregulated and downregulated respectively across all time points. Gene expression cluster analysis identified the activation of the photosynthesis pathway and photosynthesis related genes during this transition. miRNA expression analysis identified a steady increase in expression of two miRNAs, miR159a and miR159b during the transition from dormancy to budburst. Our analysis suggests that these two miRNAs promote budburst by repressing the expression of auxin responsive genes. Finally, a continued increase in CG methylation levels was observed during the dormancy to budburst transition. Differential methylation analysis, considering dormant buds as the control stage, yielded 6354 differentially methylated regions across the genome. Two glucosidase genes exhibited increases in promoter methylation and a corresponding decrease in gene expression in our analysis. This study provides a multi-omics view of grapevine bud transition from dormancy to bud burst and unveils the interacting genetic and epigenetic networks regulating this process.

## Introduction

Grapevine (*Vitis vinifera*) is an economically important fruit crop, well adapted to grow in wide range of climatic conditions (Sargolzaei et al., 2021), which is mainly grown for wine production. Throughout the 2021/2022 season, global grapevine production amounted to 25.62 million metric tons (M. Shahbandeh, 2022). In the United States alone, which ranks fourth in wine production by volume (Stevenson, 2005), wine, grapes, and grape products contributed $276 billion to the economy in 2022 (Wine America, 2022). Grapevine importance goes beyond economics. Since its genome sequence was released, grapevine has become a model crop for fruit genetics studies and is one of the most widely studied fruit crops among non-climacteric fruit crops (Velasco et al., 2007; Zombardo et al., 2020).

During unfavourable conditions, plants temporarily suspend growth using a set of molecular, physiological, and morphological processes, collectively called dormancy. Dormancy is common survival strategy of plants, especially in temperate woody perennials. Based on the source of environmental signals that induce dormancy, it is categorized into three classes: paradormancy (PD), endodormancy (ED), and ecodormancy (ECD) (Liu and Sherif, 2019a). PD, also called summer dormancy, is the temporary dormancy which precedes ED in woody plants. PD is mainly controlled by dominant bud growth i.e., apical dominance (Cline and Deppong, 1999). In ED, low temperature stimulates internal bud signals, which enable buds to become tolerant to low temperature. ED restricts new growth until the fulfillment of the chilling requirement (Yang et al., 2021). ECD is last stage of dormancy where buds resume new growth after getting the correct environmental signal. This resumption of new growth after ECD is often called budburst or bud break (Camargo Alvarez et al., 2018).

Bud dormancy and budburst are important agronomical traits as they control vegetative growth. Budburst is the stage when dormant plants initiate new growth from their dormant buds, signalling the end of dormancy, and the beginning of the new growing season (Lang, 1987). Timing of budburst helps to synchronize flowering, fruit set, and harvest. Much of the supply of grapevine fruits is reliant on the successful timing of budburst (Beauvieux et al., 2018). Phytohormones, xylem pressure, tissue oxygen status, callose deposition, photoperiod, and soil temperature are already reported to be involved in this phase transition (Meitha et al., 2015; Signorelli et al., 2018a; Signorelli et al., 2020).

Dormancy to budburst phase transition is a complex process, with many physiological, molecular, and biochemical changes occurring during the process. Little is known about the molecular processes underlying this transition in grapevine. Previous reports using next-generation sequencing in sweet cherry predicted a complex array of signaling pathways, including cold responsive genes, abscisic acid (ABA)-responsive genes and DORMANCY ASSOCIATED MADS-box genes involved in bud dormancy and budburst (Vimont et al., 2019). Transcriptome analysis of hazelnut buds during the ECD and bud burst stages has shown that the phenylpropanoid and phytohormone biosynthesis pathway genes are significantly enriched during budburst, indicating that such pathways play critical roles in regulating bud burst (Kavas et al., 2019). This study further reported that bud burst involves the interaction between plant growth inhibiting substances such as, abscisic acid and other plant hormones including auxin, gibberellins, and ethylene (Kavas et al., 2019). A recent study in grapevine (cultivar Merlot) by Velappan et al., identified the influence of ABA-signaling in the bud dormancy (winter bud) to budburst (spring bud) transition. This research has also emphasized the significance of tissue oxygen levels and the metabolic activity of buds during this transitional phase (Velappan et al., 2022).

Our research study was planned to investigate the molecular processes involved in bud ecodormancy to budburst (partial bud opening) transition in grapevine (*Vitis vinifera* L. cv. Cabernet Sauvignon). RNA sequencing, whole-genome bisulphite sequencing and small RNA sequencing were performed to study changes in protein coding and small RNA gene expression, and DNA methylation at five time points during budburst. Findings of our analysis will contribute to an understanding how such molecular processes interact to regulate the transition between bud dormancy and bud burst.

## Materials and methods

### Plant material

Cabernet Sauvignon plants were vegetatively propagated using callused cuttings (CC) as described previously (Grigg et al., 2018), from approximately 60 cm canes collected from endodormant vines planted at the UC-Davis experimental vineyard during the Fall of 2019 and stored at 4°C for three months to break endodormancy. In May 2020, 15 canes were randomly selected and trimmed to sections containing 3 nodes (approximately 20cm), the lower end was dipped in rooting gel (IBA, 3mg/litre; clonex growth technology) for 2 seconds and placed in 24 cm pots containing 1:1 prolite-promix mixture and placed in a mist bed. During the experiment, soil temperature and soil moisture were monitored using soil sensor (WatchDog 1000 Series Micro Stations - External Sensors-3685WD1). The soil temperature throughout the experiment fluctuated between 27-33°C, whereas soil moisture was 3-5%.

The 15 CC were randomly assigned to 5 groups, with each group containing 3 replicates. Each group was used for plant material collection at one of the five time points selected for this study. In all cases, only the top end bud was collected, and plant material was collected only once from each CC at the same time of the day (11am in the morning). The initial time point, ST0 hereafter, was defined as the day canes were taken from cold storage, so buds were sampled before cuttings were treated with IBA and planted. Such resting buds prior to planting were considered as ecodormant and considered the control to all other time points. ST1 to ST4 time points were determined according to the modified Eichhorn-Lorenz System (E-L) (Coombe, 1995). The number of days since ecodormant canes were taken from cold storage (i.e., DSCS) was recorded for each time point. ST1 (5 DSCS): Swelling observed on the top bud; ST2 (9 DSCS): Bud enlargement and bud scales opening; ST3 (11 DSCS): bud turn woolly; ST4 (17 DSCS): Leaf green tip visible, first clear indication of budburst (Figure 1). Bud samples were immediately frozen in liquid nitrogen and stored at −80°C until nucleic acid extraction.

**Figure 1:**
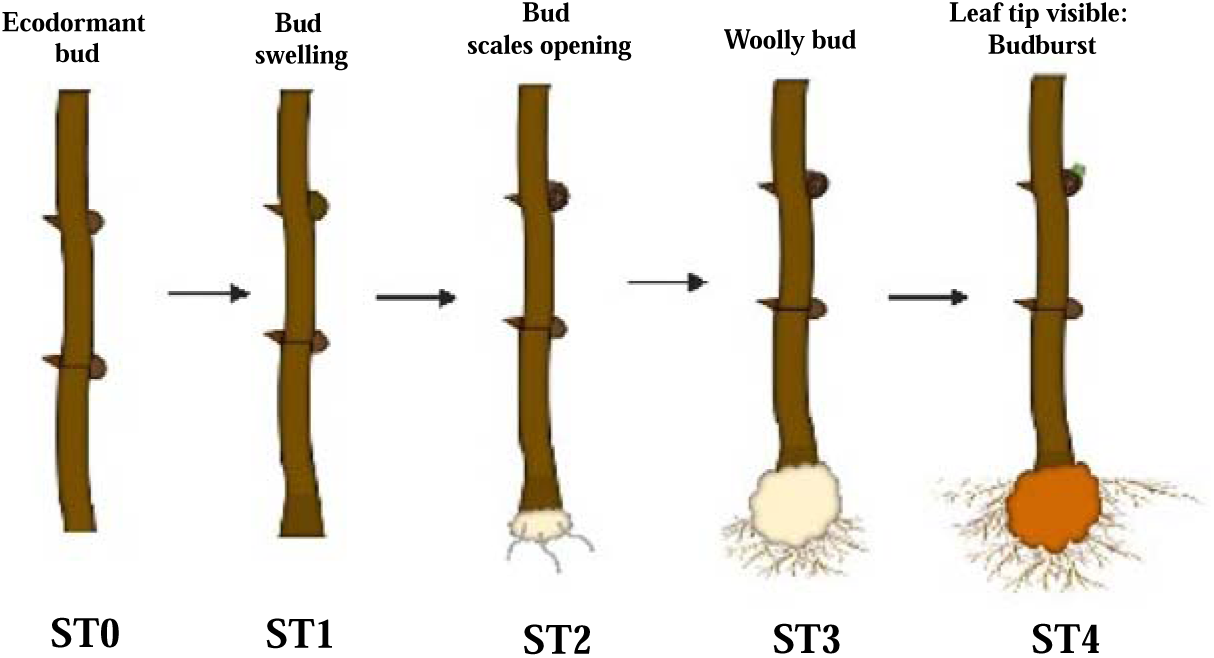
Developmental stages of budburst in grapevine. ST0-Ecodormant bud, ST1-Bud swelling, ST2: Bud scales opening, ST3: Woolly bud, ST4: Budburst. [According to the modified Eichhorn-Lorenz System (E-L)]

### Nucleic acids extraction

MPbio Fast DNA & RNA extraction kit was used for DNA and RNA from collected buds following the manufacturer’s instructions with a minor modification, i.e., the volume of 1 M DTT was also increased to 170% of the extraction kit specified amount. DNA and RNA were quantified, and quality checked using the High Sensitivity DNA or RNA Kits for Fragment Analyzer (Agilent Technologies). Only RNA samples with a RQN above 7 were kept for further use. All extracted samples were kept at −80°C until further use.

### Gene expression analysis

Total RNA extracted from bud samples was used for the expression analysis of protein coding (PCG) and long non-coding RNA genes.

Sequencing libraries for PCG expression analysis were prepared by NovoGene USA. In brief, mRNA from total RNA was enriched using oligo (dT) beads. cDNA was synthesized from the purified mRNA using random hexamer primers, after which second-strand synthesis buffer (Illumina) that included dNTPs, RNase H and DNA polymerase I added to initiate the second-strand synthesis. The double-stranded cDNA library was subjected to size selection (150 bp) and PCR enriched. Libraries were sequenced using a NovaSeq 6000 (Illumina) (2X150bp) paired end run.

Raw sequencing reads were subjected to quality control by FastQC (https://www.bioinformatics.babraham.ac.uk/projects/fastqc/). Sequencing adaptors and poor-quality reads were filtered using trimmomatic (Bolger et al., 2014).After series QC analysis, RNA-Seq reads were mapped to the grapevine reference genome PN40024.V2 (Jaillon et al., 2007) using HISAT2 default parameters (Kim et al., 2019). The python framework HTSeq (version 0.6.1) (Anders et al., 2015) was used to count the number of reads mapped to each gene. Individual gene expression levels were calculated in the form of Transcripts Per Million (TPM) for all 15 samples from their binary sequence alignment files (BAM) using the TPM calculator package (Alvarez et al., 2019). Gene expression results were used for the identification of clusters using k mean clustering (Ge et al., 2018a)and *clust* v1.8.4. (Abu-Jamous and Kelly, 2018). Principal component analysis of all 15 samples was performed using PGSEA package in iDEP.96. The count normalisation i.e., count per million (CPM) and differential gene expression were performed using DESeq2. Genes showing a fold change in expression ≥_J2 or ≤_J−2 and an FDR-corrected p values < 0.01 were deemed as differentially expressed between ST0 and ST1-4 samples. Pathway enrichment analysis was performed using KEGG pathway analysis tool using plugins of iDEP.96 web application (Ge et al., 2018b).

Small RNA libraries were generated in house using GenXPro low-input small RNA library preparation kit (GenXPro GmbH, Frankfurt, Germany) following manufacturer’s instructions. Individually indexed libraries were equimolarly pooled and sequenced using an Illumina HiSeq 150bp PE run. Quality control and adaptor trimming of raw small RNA reads were conducted using FastQC and Cutadapt respectively. miRDeep2 pipeline was used to identify and quantify known microRNAs (Friedländer et al., 2012). The target genes of miRNAs were searched using online web server psRNATarget (Dai et al., 2018).

### Whole-genome bisulphite sequencing (WGBS)

WGBS libraries were prepared by NovoGene USA (California). In brief, after a quality check, genomic DNA was spiked with unmethylated Lambda DNA and fragmented to 200-400 bp was performed using a Covaris S220 focussed ultrasonicator. Next, libraries were prepared using Accel-NGS Methyl-Seq DNA Library Kit for Illumina, (Swift biosciences) following the manufacturer’s instructions. Libraries were equimolarly pooled and sequenced using Nova Seq 6000, using a paired end (150bp) run. After sequencing, adaptor sequences, and low-quality reads were removed using Adapter Removal V2 software.

The bisulphite-converted version of grapevine reference genome (PN40024) was performed using Bismarck. Sequencing read mapping and methylation calling were performed using Bowtie2 and Bismark respectively (Krueger and Andrews, 2011; Langmead and Salzberg, 2012). Samtools (http://www.htslib.org/doc/samtools-reference.html) with --coverage and -- depth option was used to calculate sequencing coverage and depth. Differentially methylated regions (DMRs) were identified using R package methylKit (Akalin et al., 2012). The tilling window for DMRs were set at 1000kb. The regions showing difference in methylation ≥ 25% or ≤ −25% and q-value ≤ 0.01 were considered as significant DMRs.

## Results

### RNA Sequencing

Transcriptome sequencing yielded an average of 46 million reads per sample after quality filtering. The average percentage of mappable reads per sample after de-multiplexing was 90.26%, ranging from 84.33-92.85% (Supplementary file S1A). Analysis of mapping results yielded sequencing data for 26,678 genes out of the 30,661 genes annotated for this version of the *V. vinifera* genome (Supplementary file S2).

Principal component analysis (PCA) (Fig. 2A) grouped samples by sampling time point, except for one of the replicates from ST2. Samples were further grouped into three differentiated clusters. PC1 (33% of total variability) separated the cluster containing ecodormant samples (ST0) from two clusters containing ST1-ST4 samples, while PC2 (18% of total variability) separated ST1 and ST2 samples from ST3 and ST4 samples. The cluster occupying the top-right quadrant contained samples in the early stages of budbreak, i.e., all three samples from time point ST1 and two out of three samples from ST2. The cluster occupying the bottom-right quadrant contained samples in the late stages of budbreak, i.e., all three samples from time point ST3 and ST4 plus the remaining sample from ST2 (Fig. 2A).

**Figure 2.**
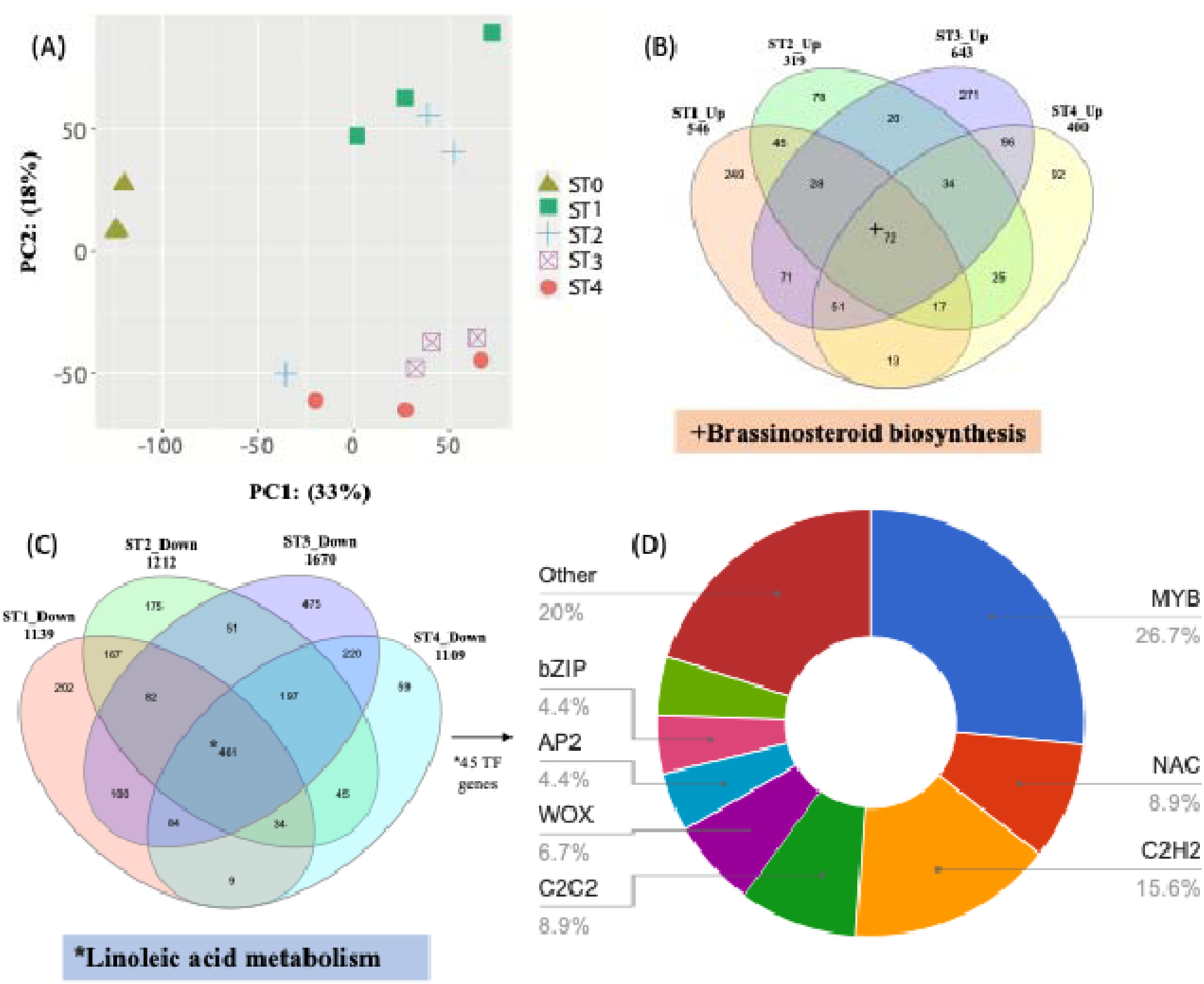
Global gene expression differences during the transition from ecodormancy to budburst in grapevine: (A) Principal component analysis of gene expression patterns bud developmental stages (ST0-ST4). ST0: Ecodormant buds, ST1: Bud swelling. ST2: Bud enlargement and bud scales opening ST2: Bud enlargement and tiny green tip appearance and ST4: Partial budburst. (B-C). Venn diagrams depicting differentially expressed genes between ST0 buds and the rest of the stages of budburst. (B) Up-regulated genes and (C) Down-regulated genes. (D) Transcription factor encoding gene families down-regulated in all budburst stages.

Differential gene expression analysis considering ST0 as a control yielded in total 3512 unique differentially expressed protein coding genes. Of these, 11 DEGs were upregulated in one or more sampling times and downregulated in the rest. Of the remaining 3501 DEGs, 1151 were upregulated and 2350 downregulated in one or more time point, 1590 genes were unique DEGs to a single time point, and 1444 and 467 were down or upregulated in two or more time points respectively. Finally, 72 and 461 DEGs were found to be up-regulated or down-regulated respectively in all time points (Figure 2B-C) (See Supplementary Fig. 1A and Supplementary File S3A-D for DEGs identified for each of the four comparisons and Supplementary File S3E for DEGs shared by more than one time point). Pathway analysis of up-regulated or down-regulated genes during all time points, showed that the former encodes for proteins involved in the brassinosteroid biosynthesis pathway, while the later encode for proteins involved in the linoleic acid metabolism pathway (Figure 2B-C). Further analysis indicates that the MYB transcription factor family contains the largest number of transcription factor genes among the commonly downregulated genes (Figure 2D). Differential expression of 142 non-coding RNAs (116 upregulated and 26 downregulated) was also observed. Non-coding RNAs were mainly snoRNAs, snRNAs and SRP-RNAs (Supplementary Fig. 1B, Supplementary File S4A-D).

### Gene Cluster Analysis

TPM calculated from BAM files were used for gene expression clustering analyses (Supplementary file S2). K-mean clustering analysis generated four gene clusters (Clusters A-D hereafter) (Figure 3; Supplementary File S5), showing distinct expression pattern during the time course of bud break. Cluster A contained 735 genes associated with cell wall biogenesis and plant hormone signalling, which exhibited higher transcript abundance during bud ecodormancy (ST0) and a subsequent abrupt decrease in the following stages of budbreak. Cluster B contained 340 genes which expression increased in the earlier stages of budbreak (ST1-2) followed by a decrease in expression in later stages (ST3-4) and which function is linked to response to oxidative stress and temperature stimulus. Genes in Cluster C (306), on the contrary showed an increase in expression on the later stages of budbreak (ST3-4) and their function is associated to transport and flavonoid biosynthesis. Finally, Cluster D contained 619 genes, which had minimal transcript accumulation during bud ecodormancy (ST0) followed by a steady increase in expression reaching a maximum transcript accumulation during bud burst (ST4). Functional annotation of cluster D discovered activation of photosynthesis and photosynthesis related genes (Figure 3).

**Figure 3:**
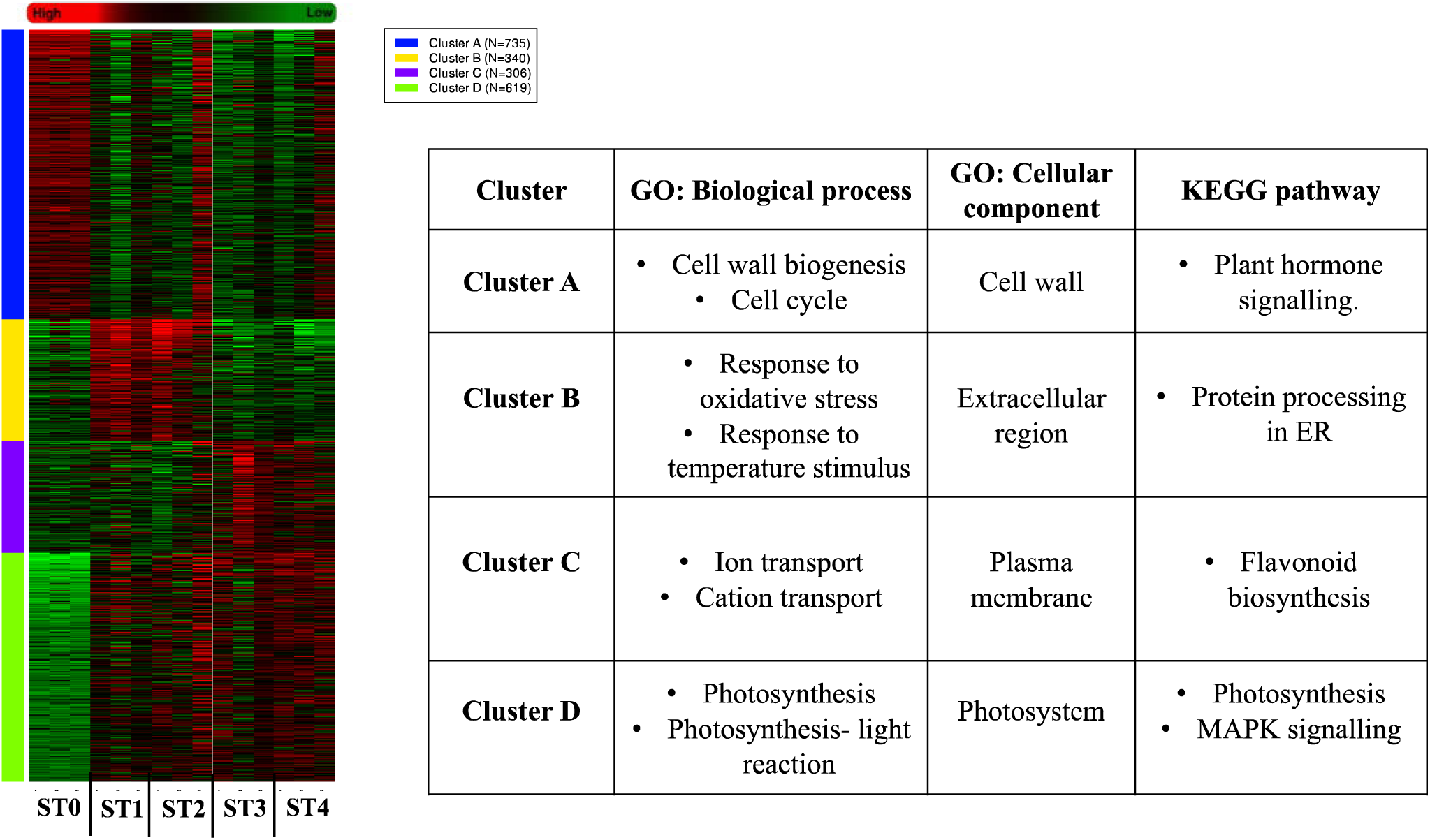
Gene expression cluster analysis during grapevine bud transition from ecodormancy to budburst: Heat map shows the gene expression levels of 2000 genes grouped into four clusters during the bud transition from ecodormancy to budburst (ST0: Ecodormant buds, ST1: Bud swelling. ST2: Bud enlargement and bud scales opening ST2: Bud enlargement and tiny green tip appearance and ST4: Partial budburst). Table insert shows the gene ontology and KEGG pathway analysis results obtained for each cluster.

Gene clustering analysis using ‘clust’ identified thirteen clusters (C0 to C12) containing 14012 genes (Supplementary Fig. 2; Supplementary File S6). Of these, clusters C0 and C7, which contained the largest number of genes (C0:3210 genes, and C7:3399 genes) and showed a gradual increase (C0) or decrease (C7) in gene expression during the bud dormancy to bud burst transition, were selected for more detailed analysis. Analysis of gene functions and pathways associated with these two clusters, identified that cluster C0 (which expression is positively associated with the transition from bud dormancy to budburst) was significantly associated with photosynthesis, especially with genes associated with photosystem I and photosystem II protein complexes (Figure 4). While, C7 main associated function was ribosome, (Supplementary Fig. 3).

**Figure 4:**
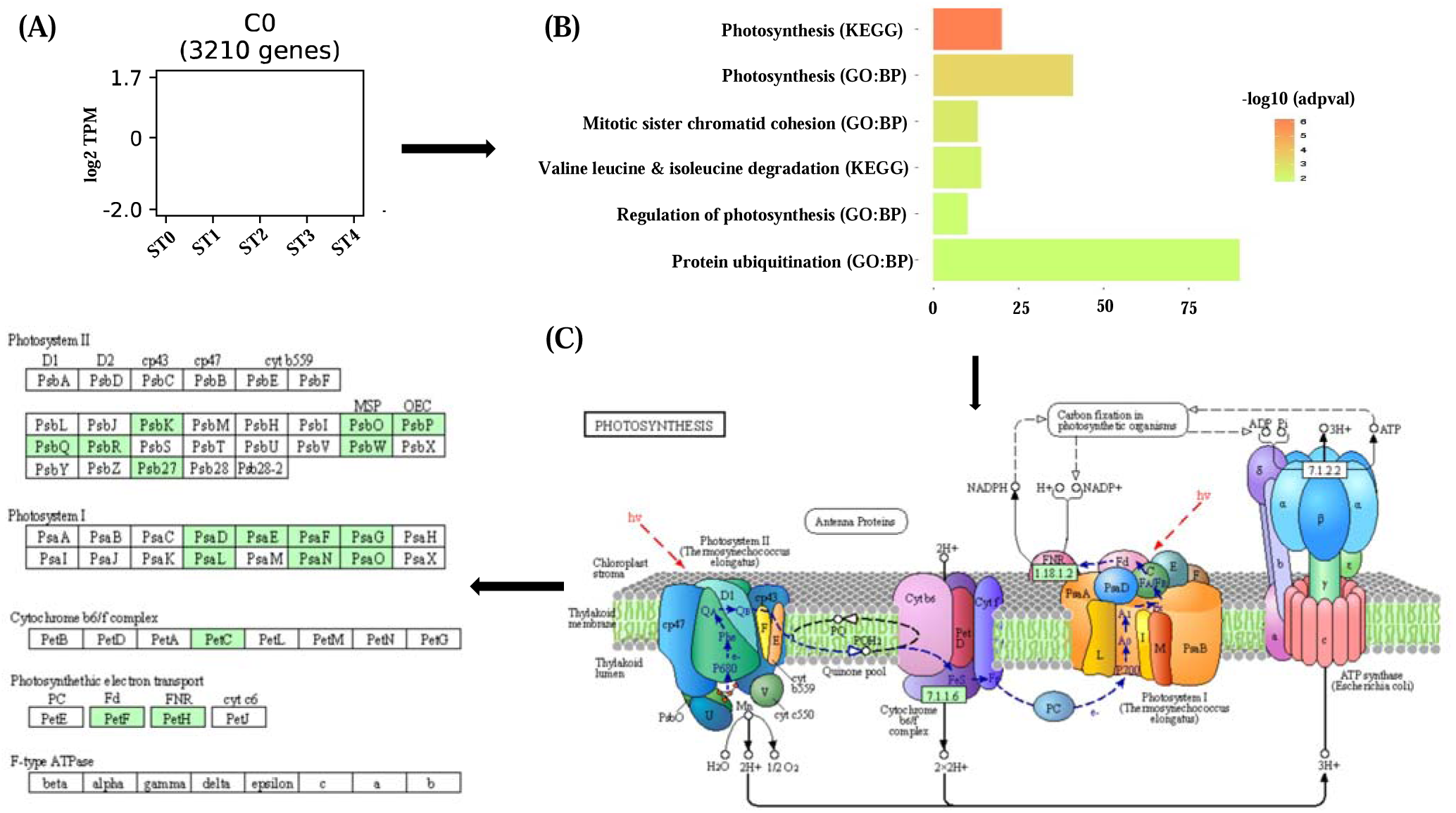
Identification of genes positively associated with bud transition from ecodormancy to budburst. A) Gene expression pattern for genes in cluster C0 (3210) showing a positive association with the ecodormancy (ST0) to budburst (ST4) transition. B) Gene ontology and KEGG pathway analysis of C0 cluster genes. C) Identification of genes associated with photosynthesis present in cluster C0. Right panel shows the photosynthesis four major membrane protein complexes (photosystem I, cytochrome b6/f, photosystem II, and ATP synthase). Left panel shows the genes identified in cluster C0 for each of the complexes, highlighted in green.

### micro-RNA analysis

The total small RNA sequencing raw read counts varied between 7 and 33 million reads, with average of 23 million reads per sample (Supplementary file S1B). After quality check and filtering, 83% raw reads were processed for miRNA identification. A total of 577 miRNA species were identified from our sequencing results (Supplementary File S7). Of these, only miRNA159a and miRNA159b showed sufficient sequencing depth for further analysis. Both miRNA159a and miRNA159b exhibited a steady increase in expression from the bud ecodormancy (ST0) to the bud burst (ST4) transition (Figure 5; Supplementary File S7). Search for their potential target genes in cluster C7 (3399 genes) yielded a total of 136 potential targets (Supplementary file S8). Biological function of these target genes identified plant hormone signalling pathways as the most significant pathway we identified increased transcript abundance of auxin induced genes.

**Figure 5:**
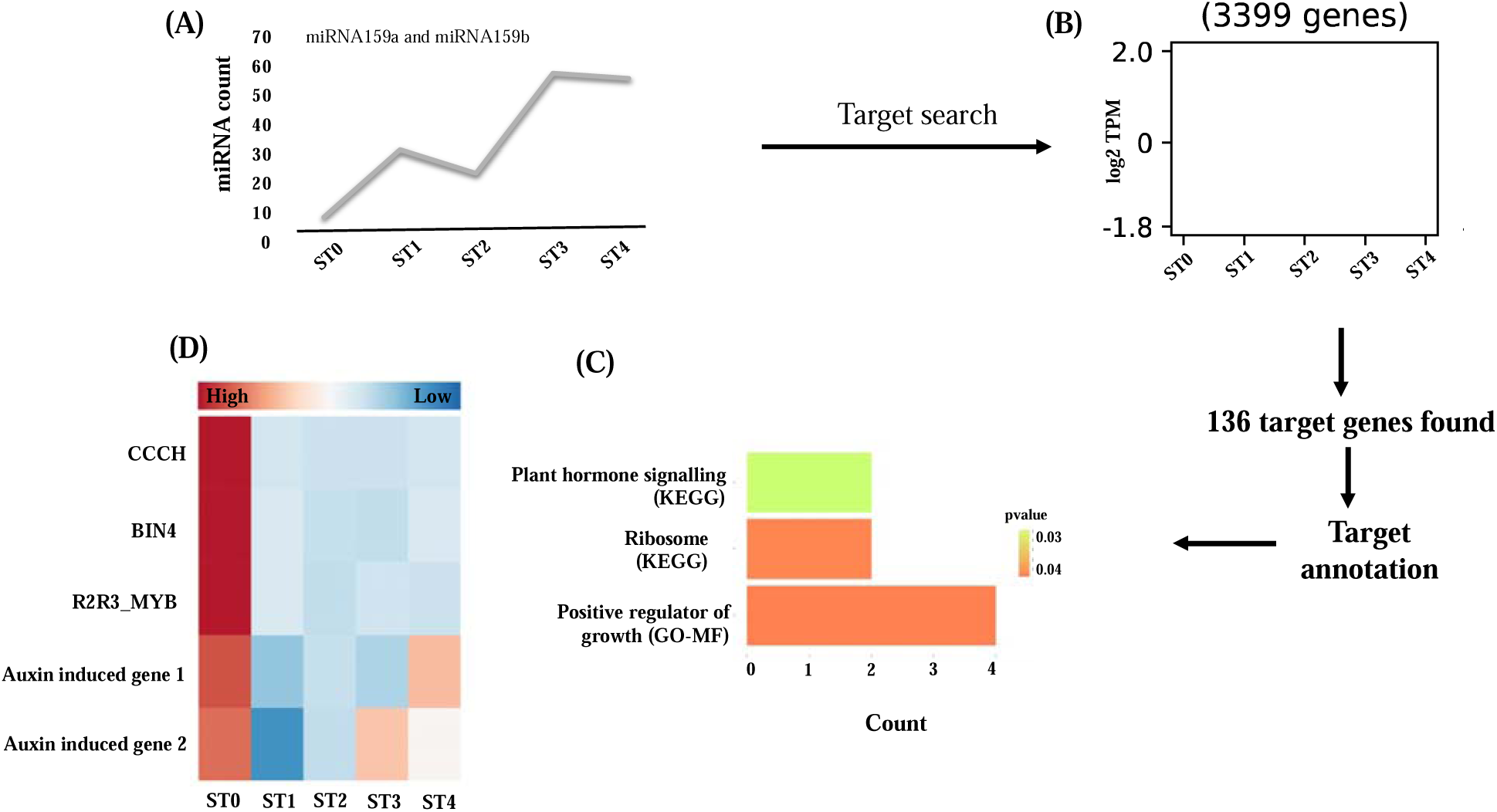
miRNA159 and miRNA159b acts as a positive regulator of budburst. (A). Expression levels (counts) of miR159a and miR159b during grapevine’s bud dormancy to budburst transition. (B) Identification of potential target genes miR159a and miR159b within genes alocated to cluster 12 using clust, which show a decrease in expression during this transition. (C) Significant target genes encode plant hormone signalling pathway. (D) Heat map depicting expression of target genes including auxin induced genes.

### DNA methylation

Whole genome methylome sequencing yielded a total of 1.49 billion reads. Bisulfite conversion efficiency showed on average 99.72% unmethylated cytosines converted to uracils. Average genome coverage was 80.2%, ranging from 78.66-82.01%, and the average sequencing depth reached was 9%, ranging from 7.1-12.48 % (Supplementary file S1C). The average percentage of mappable reads per sample after de-multiplexing was 30.43%, ranging from 25-32.90%. The average percentage of methylated cytosines (mCs) varied between 47.40% and 51.20% mCs at CG sites, 20.40-22.20% at CHGs, and 4.3-6.7% at CHHs (H=A, T or C). Analysis of global levels of DNA methylation showed a significant increase in DNA methylation in the CG and CHG contexts in the transition from eco-dormant to breaking buds (Percentage of methylated CGs and CHGs at ST0 = 47.96 and 20.9% respectively and at ST4 = 50.5 and 21.9%), while the CHH context did not show any trend (Figure 6A).

**Figure 6:**
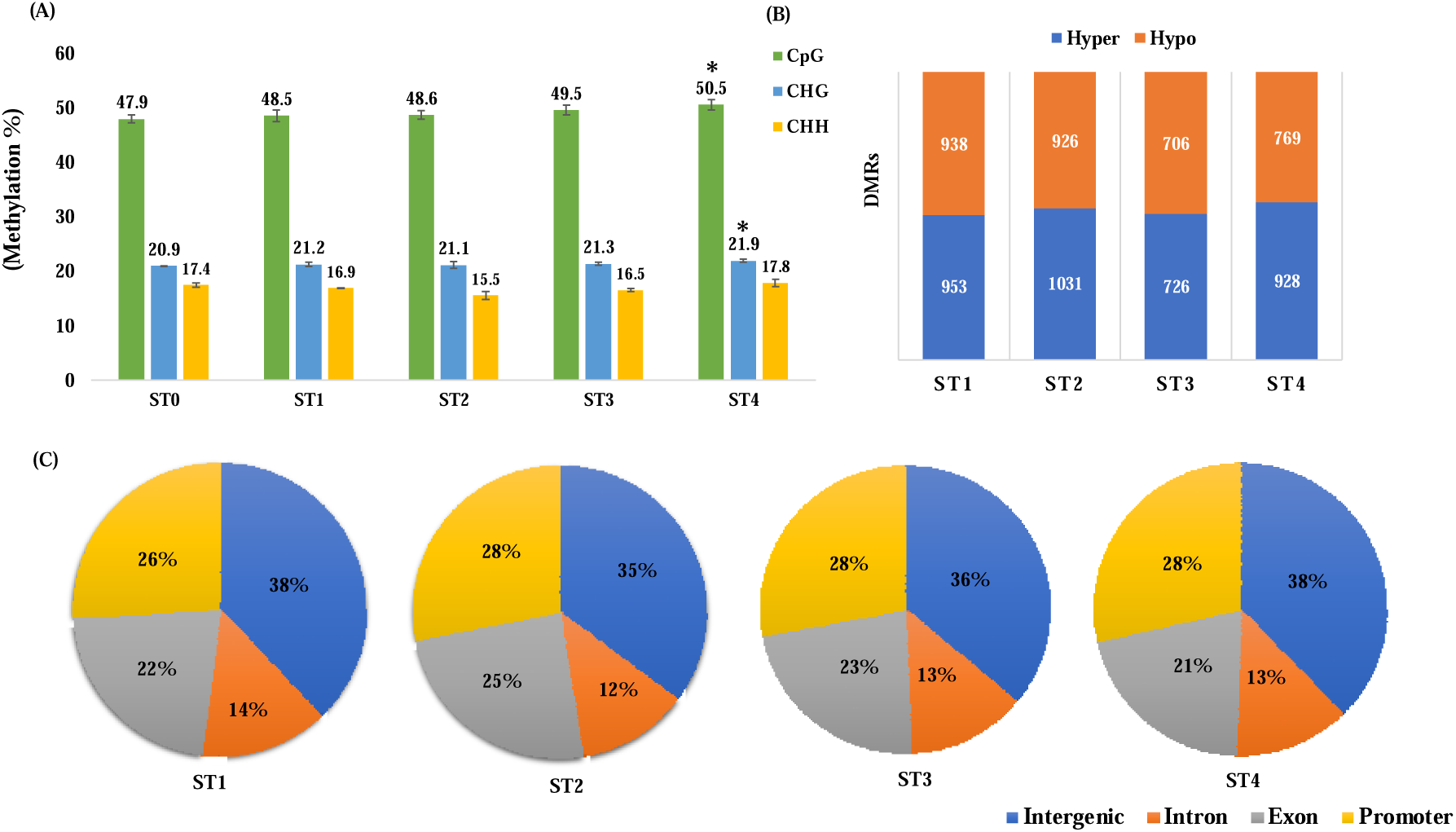
DNA methylation of grapevine buds during different bud developmental stages. (A) Methylation level in bud in the CpG, CHG, and CHH sequence contexts. Data are means ± SD of three bud samples. *: P < 0.05 vs. “ST0” in student t-test. Significant increase in DNA methylation level (CG and CHG) between eco-dormant bud and budburst stage were observed. (B) Differentially methylated regions across all stages considering ST0 as a control stage. (C) Pie charts showing the percentage methylation in intergenic regions, Intron, Exon, and Promoter.

Differential methylation analysis considering ST0 as the control stage and ST1, ST2, ST2 and ST4 as treatments yielded 6354 differentially methylated regions (DMRs), among them 3078 regions were hypomethylated (hyper-DMR) and 3276 were hypomethylated (hypo-DMR) (Figure 6B and Supplementary file S9A-D). Annotation and regional analysis exhibited DMRs were higher in intergenic regions (35 to 38%) compared to promoters (26% to 28%), exons (21 to 25%) and introns (12 to 14%) (Figure 6C).

### Gene expression and DNA methylation integration

For methylation and gene-expression integration analysis, we selected differentially expressed genes and differently methylated regions of budburst stage (ST4) compared to control (ST0). Integration of differentially methylated regions with differentially expressed genes discovered 27 genes (8 promoters and 19 gene bodies) that show hypermethylation and downregulation, 9 genes 3 promoters and 6 gene bodies) that show hyper-methylation and up-regulation, 26 genes (3 promoters and 23 gene bodies) exhibited hypo-methylation and down-regulation, and 7 genes (3 promoters and 4 gene bodies) exhibited hypo-methylation and up-regulation. *Glucosidase 12*, and genes involved in ABA response, and photosystem A/B genes showed hyper-methylation in promoter region and downregulation in gene expression. A gene encoding a *pyrophosphokinase* and two uncharacterized genes exhibited hypomethylation in promoter region and down-regulation in gene expression. In case of gene body methylation, genes like members of the families of sucrose synthase, kinesin, ethylene responsive, photosystem-I subunit, kinase and so on exhibited hypo-methylation in the gene body and down-regulation in gene expression. Genes encoding epimerase and AAA-ATPase showed hyper-methylation in the gene body and up-regulation (Figure 7).

**Figure 7:**
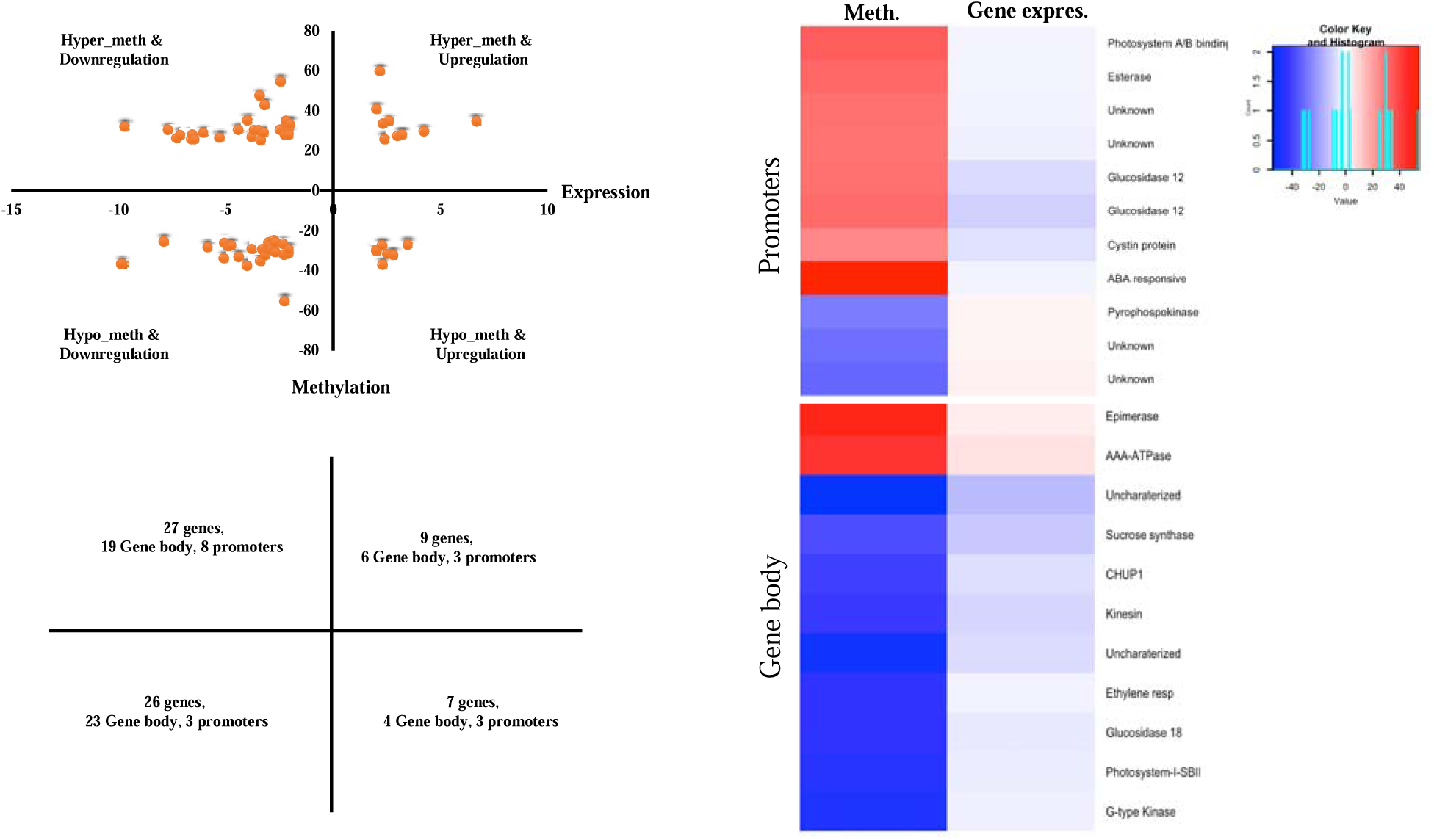
(A) Scatterplot of differentially methylated genes with differential gene expression between ST0 and ST4. (B). Heat map showing methylation and gene expression of selected promoters and gene body which shows perfect negative and positive correlation.

## Discussion

Budburst following dormancy is complex and a multifactor driven phenomenon in perennial plants. In grapevine, the molecular aspects of bud dormancy and budburst have been examined using a transcriptome approach (Khalil-Ur-Rehman et al., 2017; Shangguan et al., 2020; Noronha et al., 2021; Wang et al., 2022). The current study expands on previous work in grapevine to include a description of global gene and small RNA expression and DNA methylation pattern from same tissues to study the programmed mechanism during bud ecodormancy to bud burst transition.

The PCA analysis using transcriptome datasets represented gradual development in buds from bud dormancy to bud burst. For tree species, photoperiod, air temperature and relative water content are key drivers of budburst (Kleinknecht et al., 2015). The resting or dormant buds contains photosynthetically inactive proplastids. During the transition of these dormant buds to budburst these become functional and photosynthesis related genes get activated (Guzicka et al., 2018; Signorelli et al., 2018b). In our analysis, we observed activation of genes associated with photosynthesis, mainly genes that encode photosystem I and photosystem II components. Photosystem I and photosystem II are protein complexes that contain the pigments necessary to harvest light and use light energy to catalyse photosynthesis (Caffarri et al., 2014). Our gene clustering analyses identified photosynthesis as a significant pathway altered during budburst which gets activated during bud ecodormancy to bud burst transition. Expression of 14 different PSI and PSII related genes were found in KEGG pathway analysis. Two of the photosystems I and photosystem A/B genes showed promoter and gene body methylation and changes in their expression, suggesting that epigenetic regulation might play an important role in activating photosynthesis related genes in grapevine, and that the activation of such genes would be indicative of the transition from ecodormancy to budburst. In K-mean cluster analysis, we also observed the higher expression of plasma membrane associated genes in ST3 and ST4 as compared to ecodormant stage (Figure 3). Different studies have documented the role of plasma membrane (PM) in dormancy release. Throughout the dormancy PM remains in semi-dehydrated state. After getting external and internal signals the PM gets hydrated and activated (Aue et al., 1999; Ueno et al., 2013). This activated PM is involved in bud development during budburst. C7 cluster which shows decrease in gene expression during transition encode for ribosome pathway. Since, ribosomes control mRNA translation into proteins, our findings suggest that there might be a steady decrease in protein synthesis during the bud dormancy to budburst transition.

In plants, a major function of miR159 is to enable normal growth by silencing R2R3-MYB transcription factors (Millar et al., 2019). The miR159-MYB pathway in plants has been implicated in several different functions. In our small RNA sequencing data analysis, we observed steady increase in miR159a and miR159b from bud ecodormancy to budburst transition. This is consistent with the possibility that these two miRNAs positively regulate the budburst initiation. We observed the silencing of two auxin responsive genes and R2R3-MYB transcription factor gene by these miR159s. According to previous reports, auxin promotes the bud dormancy in woody plants (Qiu et al., 2019). Our study, suggest that miR159a and miRNA159b could be silencing auxin responsive genes to overcome bud ecodormancy and eventually initiating budburst. Through our study we report, the miRNA159-Auxin responsive gene pathway as a positive regulator of budburst and negative regulator of bud dormancy in grapevine. We also observed the expression of the long encoding RNAs such as snoRNAs, snRNAs and SnoRNAs. In peach, the regulatory role of snRNAs in ecodormancy has been reported (Yu et al., 2021)

During resumption of budburst most of the bud cells undergo cell division and cell elongation (Cooke et al., 2012). These processes are mainly controlled by plant hormones. Brassinosteroids are a class of plant hormones reported to promote cell division (Peres et al., 2019). Our differential gene expression analysis discovered brassinosteroid biosynthesis as a common upregulated pathway across all stages. A study in peach has reported that an early bud-break transcription factor gene 1 is greatly influenced by brassinosteroid metabolism (Zhao et al., 2021). Taken together, brassinosteroid acts positively in promoting budburst, our RNA sequencing results well support previous hypothesis, however detailed analysis is needed to prove the function of brassinosteoid pathway in budburst. Interestingly, the linoleic acid metabolism pathway was down-regulated during the ecodormancy to budburst transition. Linoleic acid in rapeseeds represses expression of gene regulators responsible for maintaining bud dormancy, resulting in bud growth (Huang et al., 2021). Linolenic acid initiates Jasmonic acid biosynthesis in plants (Robin Liechti and Edward E Farmer, 2006). In Japanese apricot, jasmomic acid, auxin and abscisic acid response genes are involved in seasonal bud dormancy release (Zhong et al., 2013). Linoleic acid is possibly responsible for bud dormancy release. Since our study reports down-regulation of genes with products involved in linoleic acid metabolism, there might be linoleic acid catabolism than anabolism.

MYB transcription factors (MYB-TF) genes positively regulate the budbreak in different plant species including bamboo, hazelwood and *Camellia sinensis* (Bhandawat et al., 2017; Hao et al., 2017).MYB-TFs are mostly up-regulated during budbreak in plants, so our study contradicts the previous studies by documenting the downregulation MYB-TFs across stages. However, further studies need to be performed to research how these MYB-TFs acts negative regulators of budburst in grapevine.

The observed consistent increase in CG and CHG methylation from bud ecodormancy stage (ST0) to budburst stage (ST4), suggests changes at epigenetic level during bud ecodormancy to bud burst transition. Several studies have indicated that plant hormones such as ABA, GA, auxin and cytokinin guide bud dormancy and bud burst (Horvath et al., 2008; Aksenova et al., 2013; Gillespie and Volaire, 2017). Auxin promotes bud dormancy and suppress the budburst in plants (Aloni and Peterson, 1997; Liu and Sherif, 2019b). Our integrated DNA methylation and gene expression analysis found the hypermethylation and reduced gene expression of promoter of ABA responsive gene and hypo-methylation and increased transcript abundance of ethylene responsive genes. Two of the auxin responsive genes has also been reported as targets of miR159 in this study. Previous study in oak, has discussed activation of auxin, gibberellin and brassinosteroid responses during ecodormancy to swelling bud and budburst transition (Lesur et al., 2015).

Breakdown of sugar promotes bud dormancy release and budburst (Zhang et al., 2018). This sugar signal is independent of plant hormone signalling (Hernandez et al., 2021). Sugar accumulation in dormant bud is often connected with freezing tolerance, whereas mobilization and metabolism of this reserved sugar is often related with dormancy release (de Rosa et al., 2022). In oak transcriptome analyses, stimulation of sucrose metabolism related genes was observed during ecodormancy to swelling bud transition (Lesur et al., 2015). In our DNA methylation and gene expression integrated analysis three different glucosidase genes exhibited promoter and gene body methylation and alteration in their gene expression at same time. Glucosidases are enzymes responsible for breakdown of polysaccharides into their monomers.

### Conclusion

A schematic model depicting the molecular regulators of bud ecodormancy to budburst transition grapevine is shown in Figure 8. We propose that the activation of the photosynthesi and the brassinosteoid biosynthesis pathways, MYB transcription factor, microRNAs 159a and 159b and phosphokinase genes positively regulated this transition. Whereas, Linoleic acid biosynthesis and ribosome pathways, in conjunction with Auxin induced genes and glucosidase genes found are repressors of budburst in grapevine.

**Figure 8:**
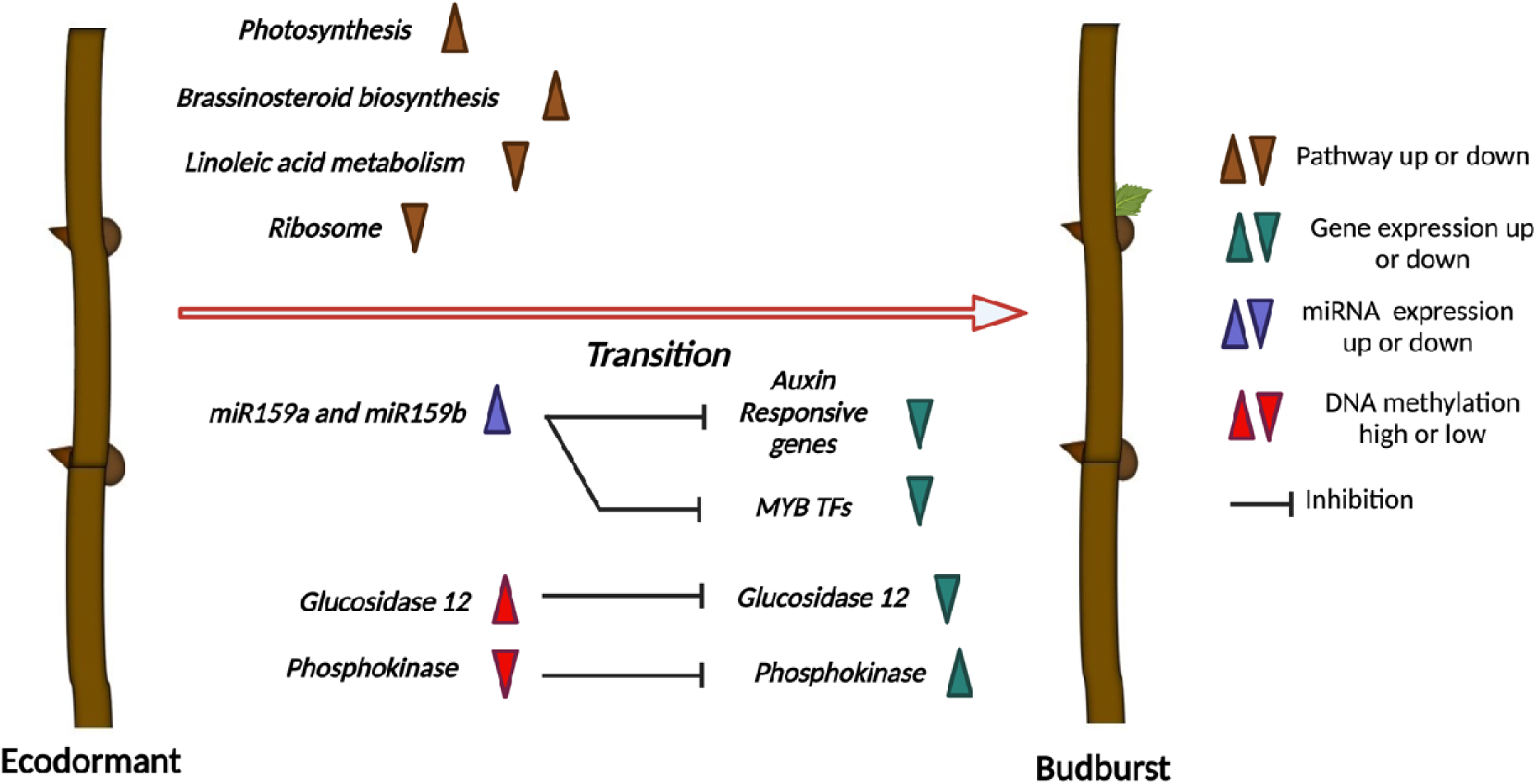
A possible model of molecular control of ecodormancy to budburst transition in grapevine. It has been shown that photosynthesis pathway, brassinosteoid biosynthesis pathway, MYB transcription factor (TFs) genes, microRNA159a, miRNA159b and phosphokinase gene acts as an activator of transition. Whereas, Linoleic acid biosynthesis pathway, ribosome pathway, Auxin induced genes and glucosidase genes are involved in repression of budburst.

## Conclusions

Our study provides several novel insights into genetic and epigenetic regulation of bud ecodormancy to budburst transition in grapevine. Multi-omics analysis reveals key changes in gene expression, microRNAs, and DNA methylation during this transition. Our results enable proposition of budburst regulating networks, such as miRNA159-Auxin responsive factors, plant hormone signalling through brassinosteroid, auxin and ABA, activation of photosynthesis and breakdown of sugar through glucosidases and so on. At molecular level, all these networks could interact and respond to external factors to regulate budburst. The novel ideas and results presented in our study, will be further explored to better understand the regulation of budburst at molecular level.

## Acknowledgements

This study was supported by the National Institute of Food and Agriculture, AFRI Competitive Grant Program Accession number 1018617, the National Institute of Food and Agriculture, United States Department of Agriculture, Hatch Program accession number 1020852.

## Author contributions

C.L. R.G., and H.S. designed the experiment. H.S., T.R., and L.A. Methodology, bioinformatics analysis, visualization, and result interpretation. H.S.-Writing manuscript draft. C.L. R.G., and S.P.-Supervision, resources, manuscript revision & editing. C.L. R.G., and S.P., Funding acquisition, Resources, Supervision.

## Data availability

The raw sequencing data were deposited to National Centre for Biotechnology Information (NCBI)-Sequence Read Archive (SRA) database. The bioproject accession number for RNA sequencing reads - PRJNA954365, small RNA sequencing reads - PRJNA955780 and Whole genome bisulfite sequencing reads-PRJNA956264.

## Supplementary files legends

**Supplementary Figure 1:** Number of upregulated and downregulated differentially expressed (A) protein coding genes and (B) non-coding RNAs during the grapevine bud ecodormancy to budburst transition in the four comparison groups (ST0 vs ST1, ST0 vs ST2, ST0 vs. ST3 and ST0 vs ST4).

**Supplementary Figure 2:** Gene expression clustering analysis using clust during grapevine’s ecodormancy to budburst transition (ST0 to ST4). In total, thirteen (C0-C12) co-expression clusters were identified.

**Supplementary Figure 3:** (A) Cluster negative regulating bud ecodormancy to bud burst transition. (B) Pathway analysis of cluster showing ribosome as significant pathway.

**Supplementary file S1:** Supplementary file S1A. RNASeq statistics for grapevine buds during their transition from ecodormancy to budburst. Three replicates (indicated in the sample ID as B1, B2, and B3) were used per sampling stage (indicated as ST0: Ecodormant buds, ST1: Bud swelling. ST2: Bud enlargement and bud scales opening ST2: Bud enlargement and tiny green tip appearance and ST4: Partial budburstof). Supplementary file S1B. Small RNASeq statistics for grapevine buds during their transition from ecodormancy to budburst. Three replicates (indicated in the sample ID as B1, B2, and B3) were used per sampling stage (indicated as ST0: Ecodormant buds, ST1: Bud swelling. ST2: Bud enlargement and bud scales opening ST2: Bud enlargement and tiny green tip appearance and ST4: Partial budburstof). Supplementary file S1C. Whole genome bisulfite sequencing statistics for grapevine buds during their transition from ecodormancy to budburst. Three replicates (indicated in the sample ID as B1, B2, and B3) were used per sampling stage (indicated as ST0: Ecodormant buds, ST1: Bud swelling. ST2: Bud enlargement and bud scales opening ST2: Bud enlargement and tiny green tip appearance and ST4: Partial budburstof).

**Supplementary file S2:** Gene expression levels (in transcripts per million (TPM) measure in grapevine buds during their transition from ecodormancy to budburst (ST0: Ecodormant buds, ST1: Bud swelling. ST2: Bud enlargement and bud scales opening ST2: Bud enlargement and tiny green tip appearance and ST4: Partial budburstof). Gene expression was measured in three replicates (B1, B2 and B3) per time point.

**Supplementary File S3A.** Identified differentially expressed protein coding genes in grapevine buds during their transition from ecodormant buds (ST0) to Bud swelling (ST1). Supplementary File S3B. Identified differentially expressed protein coding genes in grapevine buds during their transition from ecodormant buds (ST0) to Bud enlargement and bud scales opening (ST2). Supplementary File S3C. Identified differentially expressed protein coding genes in grapevine buds during their transition from ecodormant buds (ST0) to Bud enlargement and tiny green tip appearance (ST3). Supplementary File S3D. Identified differentially expressed protein coding genes in grapevine buds during their transition from ecodormant buds (ST0) to Partial budburstof (ST4). Supplementary File S2-E. Differentially expressed protein coding genes (DEGs) during grapevine transition from ecodormancy (ST0) to budburst (ST1-ST4) identified in more than one stage (ST1: Bud swelling. ST2: Bud enlargement and bud scales opening ST2: Bud enlargement and tiny green tip appearance and ST4: Partial budburst. Occurrences indicate the number of stages during budburst where a given gene was found to be differentially expressed; and differentially expressed in shows what those stages are and the directionality of the change in expression compared to ecodormant buds. Inserted figure shows the number of DEGs per stage (in parenthesis) and the number of shared DEGs between stages (in yellow boxes).

Supplementary File S4A. Identified differentially expressed non coding RNA genes in grapevine buds during their transition from ecodormant buds (ST0) to Bud swelling (ST1). Supplementary File S4B. Identified differentially expressed non coding RNA genes in grapevine buds during their transition from ecodormant buds (ST0) to Bud enlargement and bud scales opening (ST2). Supplementary File S4C. Identified differentially expressed non coding RNA genes in grapevine buds during their transition from ecodormant buds (ST0) to Bud enlargement and tiny green tip appearance (ST3). Supplementary File S4-D. Identified differentially expressed non coding RNA genes in grapevine buds during their transition from ecodormant buds (ST0) to Partial budburstof (ST4).

Supplementary file S5: Gene clusters identified by k-means clustering analysis from gene expression data of grapevine buds collected during transition from ecodormancy to budburst (ST0: Ecodormant buds, ST1: Bud swelling. ST2: Bud enlargement and bud scales opening ST2: Bud enlargement and tiny green tip appearance and ST4: Partial budburst). Three replicates per time point were used for analysis (B1-3). Columns C to Q show the log2 TPM values for all genes identified for each of the 4 clusters in each replicate.

Supplementary file S6: Genes belonging to 13 clusters (C0 to C12) identified using clustering method clust from gene expression data of grapevine buds collected during transition from ecodormancy to budburst (ST0: Ecodormant buds, ST1: Bud swelling. ST2: Bud enlargement and bud scales opening ST2: Bud enlargement and tiny green tip appearance and ST4: Partial budburst). Figure in parenthesis indicates the number of genes identified per cluster.

Supplementary file S7: Known microRNAs expression levels (in transcripts per million (TPM) measure in grapevine buds during their transition from ecodormancy to budburst (ST0: Ecodormant buds, ST1: Bud swelling. ST2: Bud enlargement and bud scales opening ST2: Bud enlargement and tiny green tip appearance and ST4: Partial budburstof). Gene expression was measured in three replicates per time point.

Supplementary file S8: Putative target genes of microRNA159a, microRNA159b identified using psRNATarget tools.

Supplementary file S9: Differentially methylated region after considering ST0 as a control stage in all bud developmental stages.

