## Supplementary figures and images for "Multi-Omics Insights into Grapevine Ecodormancy to Budburst Transition: Interplay of Gene Expression, miRNA Regulation, and DNA Methylation"

### Supplementary Figure 1

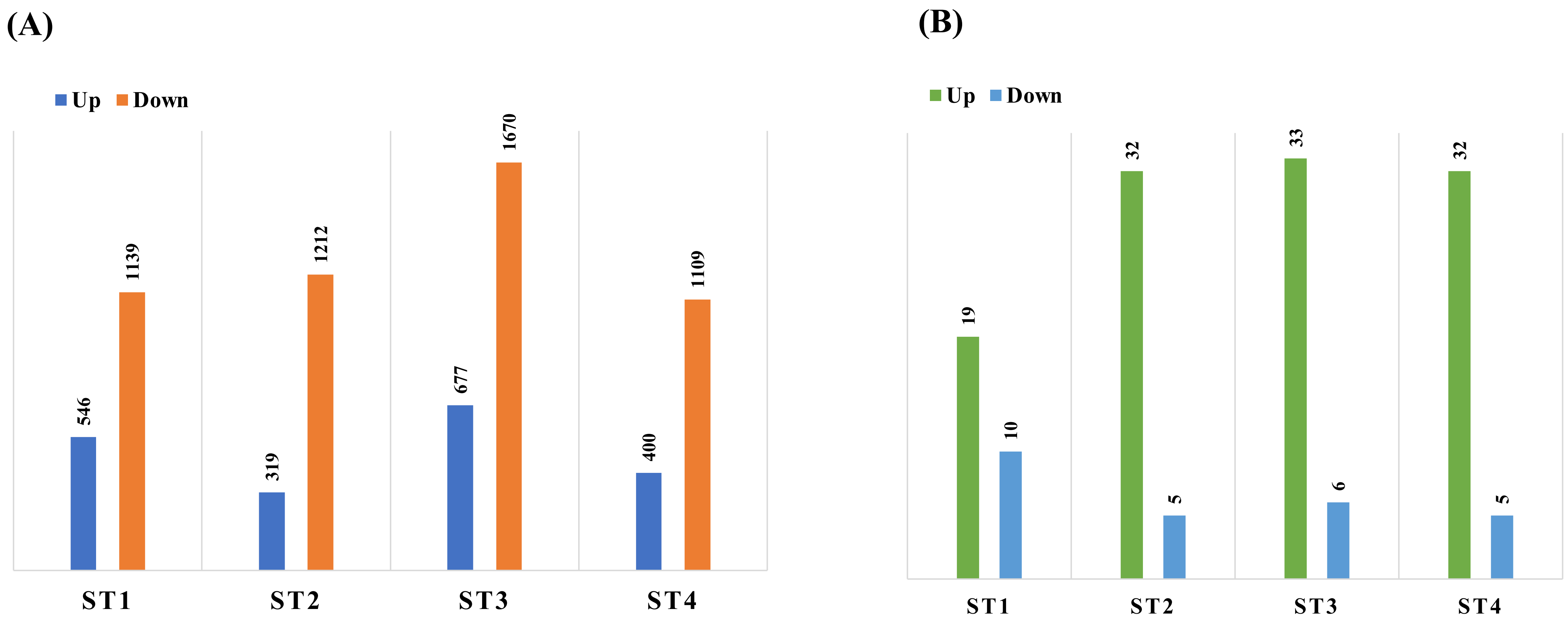

### Supplementary Figure 2

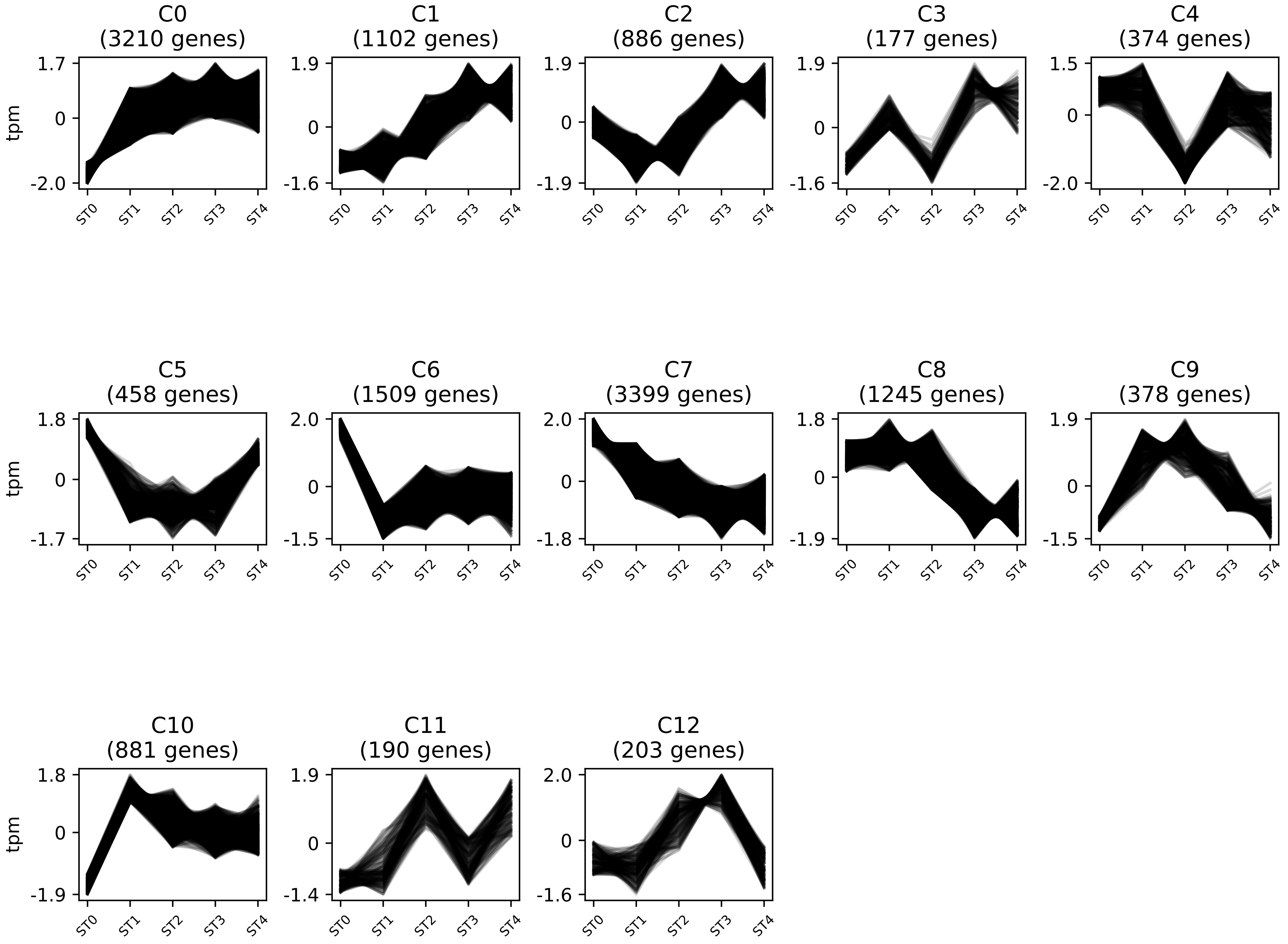

### Supplementary Figure 3

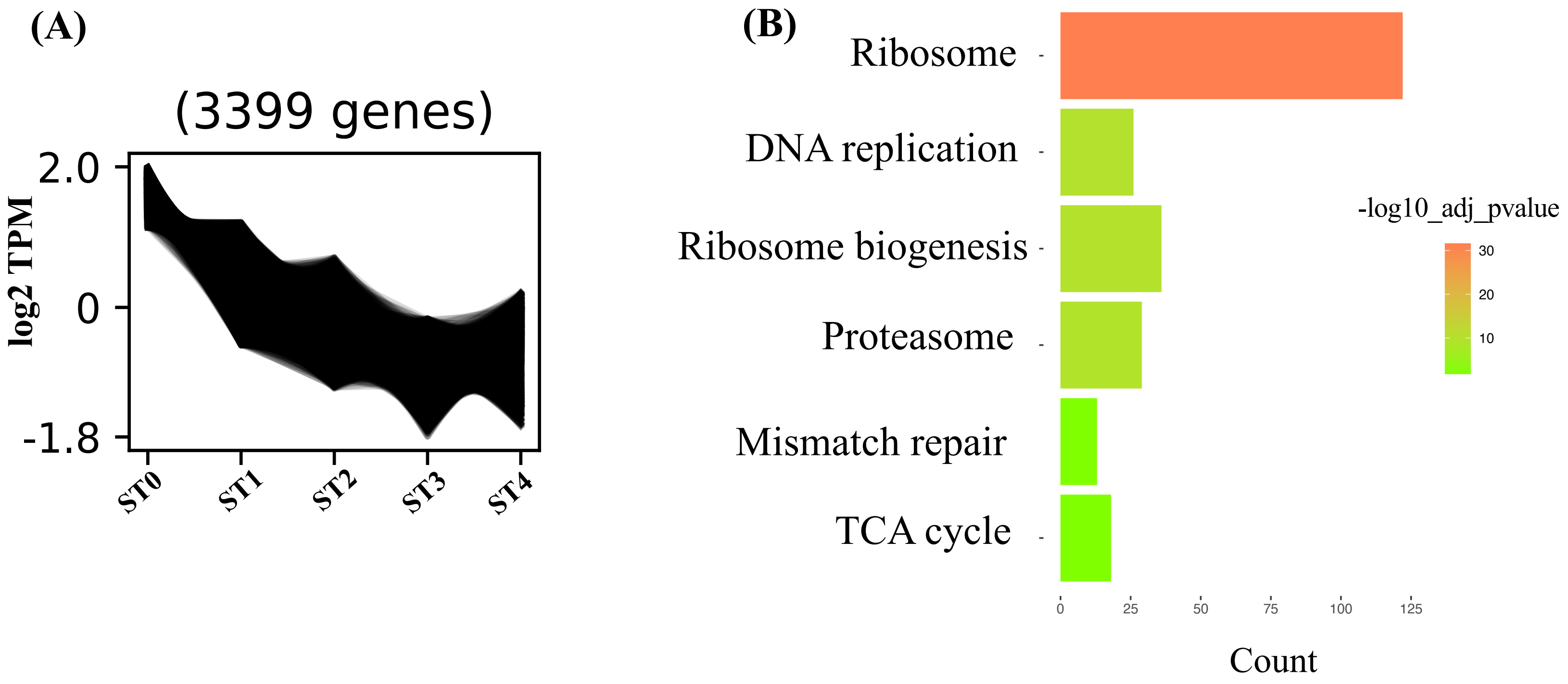
